# Leaf nitrogen does not explain changes in soybean radiation-use efficiency in vegetative and early reproductive stages

**DOI:** 10.1101/2022.03.24.485711

**Authors:** Nicolás Cafaro La Menza, Timothy J. Arkebauer, John L. Lindquist, Juan Pablo Monzon, Johannes M.H. Knops, George Graef, David Scoby, Réka Howard, Jennifer Rees, James E. Specht, Patricio Grassini

**Author notes:** Department of Health and Environmental Science, Xi’an Jiaotong-Liverpool University, Suzhou, China (Johannes M.H. Knops). Email addresses: Nicolás Cafaro La Menza, Timothy J. Arkebauer, John L. Lindquist, Juan Pablo Monzon, Johannes M.H. Knops, George Graef, David Scoby, Réka Howard, Jennifer Rees, James E. Specht, and Patricio Grassini.

## Abstract

Ontogenic changes in soybean radiation-use efficiency (RUE) have been attributed to variation in specific leaf nitrogen (SLN) based only on data collected during seed filling. We evaluated this hypothesis using data on leaf area, absorbed photosynthetically active radiation (APAR), aboveground dry matter (ADM), and plant nitrogen (N) concentration collected during the entire crop season from seven field experiments conducted in a stress-free environment. Each experiment included a full N treatment that received ample N fertilizer and a zero N treatment that relied on N fixation and soil N mineralization. We estimated RUE based on changes in ADM between sampling times and associated APAR, accounting for changes in biomass composition. The SLN and RUE exhibited different seasonal patterns: a bell-shaped pattern with a peak around the beginning of seed filling, and a convex pattern followed by an abrupt decline during late seed filling, respectively. The level of N supply influenced SLN more than RUE *via* changes in leaf N concentration, with small changes in specific leaf weight. Changes in SLN explained the decline in RUE during seed filling but failed to predict changes in RUE in earlier stages. A simple approach based on phenological stages may give more realistic estimates of RUE before seed filling, improving crop growth and yield prediction *via* crop models and remote sensing.

**Highlight:** Changes in radiation-use efficiency during soybean vegetative and early reproductive stages are not related to specific leaf nitrogen.

## 1. Introduction

Radiation use efficiency (RUE) is often defined as the amount of phytomass produced per unit of solar radiation absorbed or intercepted by the vegetation over a given period of time (Kiniry *et al*., 1989; Monteith *et al*., 1977; Sinclair and Muchow, 1999). The RUE can be used as a parameter for estimating crop productivity using remote sensing (Garbulsky *et al*., 2011), simple empirical models (Monteith, 1972), or process-based simulation models (Chapman *et al*., 1993; Jones and Kiniry, 1986; Sinclair, 1986; Villalobos *et al*., 1996). The first estimates of RUE for soybean were derived from the slope of the linear regression between cumulative aboveground dry matter (ADM) and the intercepted solar radiation during the crop cycle (Shibles and Weber, 1965). Subsequent estimates of soybean RUE have followed the same approach (Andrade, 1995; Daughtry *et al*., 1992; Muchow, 1985; Muchow *et al*., 1993b; Pengelly *et al*., 1999; Ries *et al*., 2012; Sinclair *et al*., 1987; van Roekel and Purcell, 2014). A limitation of this approach is that using cumulative values of ADM and intercepted radiation may mask changes in RUE within the crop season (Sinclair and Muchow, 1999). Indeed, the few studies that have estimated RUE for specific crop phases in soybean have detected variation in RUE during crop ontogeny. For example, Muchow (1985) observed lower RUE during seed filling, attributing the decline in RUE to a loss in photosynthetic capacity during seed filling. Similarly, other studies based on eddy covariance gas-exchange measurements have shown lower RUE early in the crop season but have not determined its underlying drivers (Dold *et al*., 2017; Gitelson and Gamon, 2015; Rochette *et al*., 1995). To summarize, this empirical evidence suggests that RUE is not constant during the crop season in soybean.

Relative to the possible causes for variation in RUE with ontogeny, Sinclair and Horie (1989) postulated that, in the absence of water stress and within the optimal temperature range, and after normalization for differences in diffuse radiation, changes in soybean RUE could be explained by changes in specific leaf nitrogen (SLN). The only empirical validation of this hypothesis was reported by Sinclair and Shiraiwa (1993) using data from field-grown soybean in Japan and Florida, USA. However, their evaluation was incomplete, as plant samples were collected only during the seed-filling phase (R5-R7)^1^ without assessing changes in RUE and SLN during vegetative and early-reproductive stages. Similarly, the range of SLN reported (1.2-1.6 g N m^-2^) was much more narrow than SLN reported for soybean in other studies, which range from *ca.* 0.7 to 2.7 g N m^-2^ leaf (Archontoulis *et al*., 2020). Sinclair and Shiraiwa (1993) also reported no influence of additional nitrogen (N) supply on RUE, which is not consistent with recent evidence of higher RUE due to the addition of N fertilizer in soybean crops grown in near-optimal conditions (Cafaro La Menza *et al*., 2020). An assessment of the effect of SLN on the RUE over the *entire* crop cycle, as influenced by N supply, is yet to be reported for soybean.

Synthesizing carbohydrates requires comparably less energy than what is needed to synthesize protein and oil (Amthor, 2010; Muchow *et al*., 1993b; Sinclair and Horie, 1989). As a result, RUE estimated based on standing dry matter is lower for crops producing harvestable organs rich in oil and/or protein (e.g., soybean, sunflower) than crops producing carbohydrate-rich end products (e.g., maize, rice, potatoes)(De Vries *et al*., 1983). For the same reason, RUE can also change within the crop season because producing oil- or protein-rich seeds is energetically more ’expensive’ than vegetative organs such as stems, petioles, roots, and leaves (Sinclair and Muchow, 1999). For example, Muchow *et al*. (1993b) estimated that seed biomass contains 1.3x higher energy content than vegetative biomass in soybean. Hence, robust assessment of seasonal changes in RUE in soybean requires accounting for differences in biomass construction costs among crop stages. Previous studies have used tabulated values for different compounds (carbohydrates, lipids, proteins) to account for differences in biomass composition during the crop season when estimating RUE (Andrade, 1995; Hall *et al*., 1995). An alternative approach consists of measuring each plant organ’s combustion heat, N concentration, and ash content (Williams *et al*., 1987). This approach allows expressing ADM in terms of glucose equivalents rather than on a dry-matter basis. Regardless of the method, we note that only a few studies have accounted for differences in construction costs among soybean organs (Amthor *et al*., 1994; Koester *et al*., 2014). Moreover, all these studies focused on plant samples collected at physiological maturity. We are not aware of any study that has accounted for seasonal changes in biomass construction costs to assess intra-seasonal variation in soybean RUE.

There is a lack of knowledge about the physiological drivers of changes in RUE during ontogeny in soybean. Here we assess the degree to which seasonal RUE is affected by changes in SLN as influenced by N supply after accounting for biomass construction costs. We used data collected from seven high-yield soybean experiments (>5 Mg ha^-1^). The database included detailed measurements of absorbed photosynthetically active radiation (APAR), SLN, ADM, and combustion heat, providing a unique opportunity to assess changes in RUE during the crop season while simultaneously investigating its underlying drivers.

## 2. Materials and Methods

### 2.1 Field experiments and nitrogen treatments

Field experiments were conducted during two soybean crop seasons (2016 and 2017) in center pivot-irrigated producer fields at four sites in the state of Nebraska, USA: Atkinson, Mead, Saronville, and Smithfield. Experiments were situated in a high yield (>4.5 Mg ha^-1^) area of each field as determined by prior years’ yield maps. The data from Atkinson in 2016 was excluded because of a severe infestation of powdery mildew (*Microsphaera diffusa*). Each of the seven site-year combinations is referred hereafter to as an ’environment’. More details about these experiments can be found elsewhere (Cafaro La Menza *et al*., 2020; Cafaro La Menza *et al*., 2017; Cafaro La Menza *et al*., 2019). These previous studies aimed to understand the influence of N supply on soybean yield, N accumulation, and whole-season RUE. Here we focus on understanding the drivers for changes in RUE *within* the season, as influenced by N supply. Across the environments and treatments, seed yield ranged from 5.3 to 6.7 Mg ha^-1^ (average: 5.8 Mg ha^-1^), while plant N accumulation ranged from 364 to 485 kg N ha^-1^. These values are in the upper range of seed yield and plant N uptake values reported in the literature for soybean, largely exceeding those reported in previous studies assessing ontogenetic changes in RUE (Dold *et al*., 2017; Gitelson and Gamon, 2015; Muchow, 1985; Rochette *et al*., 1995).

On-site portable weather stations were installed within 50-m of each experiment to track solar radiation, temperature, and precipitation throughout the growing season. In all cases, soybean was sown the year after maize. Seeds were treated with fungicide and insecticide. Seeds were not inoculated with *Rhizobia* because previous reports have shown no yield response associated with this practice in soybean crops grown in rotation with maize in fertile soils of the US-North Central region (Carciochi *et al*., 2019; De Bruin *et al*., 2010; Leggett *et al*., 2017). Crops were sown earlier than average sowing dates in the region (May 5) to ensure a high yield potential. Row spacing was 0.76 m across all environments, with seeding rates ranging from 25 to 44 seeds m^-2^. Foliar applications of herbicide, fungicide, and insecticide kept crops near-free of biotic stresses. Based on soil test results and crop nutrient requirements, pre-sowing nutrient fertilizer application ensured that soybean crops were not limited by nutrients (except for N in the zero-N treatment). Similarly, periodic water applications mitigated water stress during rain-free periods. Plant-available water in the soil remained above 50% in the upper 1.4 m during the growing season as monitored with Watermark® sensors (Irmak *et al*., 2014).

Each experiment consisted of two treatments with contrasting N supply, which served as a source of variation in SLN. There were four replicates per N treatment, which were arranged in a completely randomized design. The size of each experimental unit was 176 m^2^. The two levels of N treatment were: (a) ’zero-N’, in which the crop relied on indigenous sources of soil N supply and biological N fixation and, (b) ’full-N’, purposely designed to supply the crop with ample N from fertilizer to meet the expected seasonal crop N demand. The total applied N fertilizer in the full N treatment summed up to 870 kg N ha^−1^, which was determined based on (a) site-specific yield potential estimated using SoySim crop simulation model (Setiyono *et al*., 2010), (b) soybean requirement of 80 kg of N uptake in ADM per Mg^−1^ seed yield (Salvagiotti *et al*., 2008; Tamagno *et al*., 2017), and (c) an additional 40% of fertilizer to offset potential N losses including N volatilization and leaching. The total amount of N fertilizer was split into five applications to match the expected N accumulation pattern at key stages during the growing season (Bender *et al*., 2015; Thies *et al*., 1995). The five splits consisted of 10%, 10%, 20%, 30%, and 30% of the total N fertilizer, and applied at or near the stages of V2 (two developed trifoliolate leaves), V4 (four developed trifoliolate leaves), R1 (beginning of flowering), R3 (beginning of pod setting), and R5 (beginning of seed filling), respectively. The N fertilizer was applied as urea and broadcasted between plant rows.

### 2.2. Measurements of crop phenology, aboveground dry matter, and construction costs

The crop development was tracked weekly from emergence (VE) to physiological maturity (R7) by inspecting 10 consecutive plants in each replicate following the phenological scale developed by Fehr and Caviness (1977). A dimensionless development stage (DS) scale adapted from Lindquist *et al*. (2005) was used to allow comparisons among environments avoiding biases arising from site-specific weather, sowing date, and cultivar maturity group. The DS in each environment was calculated based on a beta function model relating crop development rate and daily mean air temperature, using appropriate cardinal temperatures for each phenological phase (Setiyono *et al*., 2007; Wang and Engel, 1998). In our DS scale, DS=0, 1, and 2 correspond to VE, R3, and R7 stages, respectively, and there is no need to account for photoperiod because the environments were located within a narrow latitudinal band (from 40.5°N to 42.6°N).

Seasonal ADM accumulation was determined by collecting plant samples weekly, from VE until R7. This sampling interval was considered adequate for assessing intra-seasonal changes in RUE (Hall *et al*., 1995; Sinclair and Muchow, 1999). Plants within a 1-m row (0.76 m^2^) in each replicate were clipped at the soil surface. Each sampling area was surrounded by two undisturbed rows receiving the same N treatment and separated by one meter from the adjacent sampling area within the same row. Plastic nets of 1-m length were placed between rows to collect abscised leaves. Each plant sample was separated into green leaves, stems plus petioles, seeds, pod walls, and senesced leaves. Standing leaves were assigned to the “green leaves” category when their green area was >50%. Otherwise, standing leaves were assigned to the “senesced leaves” category. Green leaf area index (LAI) was determined for each weekly sample from emergence to physiological maturity using an LAI-3100 area meter (LI-COR, Lincoln, NE). Every plant organ sample was oven-dried at 70 C° until reaching constant weight. Each plant organ sample was separately ground in a Wiley mill (1-mm screen mesh), after which the sample N concentration was determined with a dry combustion-based analyzer (LECO Corporation, St Joseph, MI). Accumulated ADM and N at a given sampling time was calculated as the sum of dry matter or N in leaves (including green, senesced, and abscised), stems plus petioles, seeds, and pod walls. We estimated SLN (g N m^-2^ leaf) as the ratio between accumulated N in green leaves and green LAI, while specific leaf weight (SLW, g leaf dry matter m^-2^ leaf) was calculated as the ratio between accumulated dry matter in green leaves and green LAI.

Biomass construction costs for each plant organ were calculated based on combustion heat, total nitrogen content, and mineral fraction following Williams *et al*. (1987). This method predicts the amount of glucose required to provide carbon skeletons, reductants, and ATP to synthesize a gram of plant organ following common biochemical pathways. We measured combustion heat using a Parr® 1108 oxygen bomb calorimeter separately for each plant organ (green leaves, senesced and abscised leaves, stems plus petioles, pod walls, and seeds). Thus, the dry matter of each organ was multiplied by the average construction cost estimated for the same organ at a given phenological stage and then summed up to estimate the overall ADM construction costs (ADM_CC_) at each sampling time and expressed as grams glucose equivalents per unit of sampled area (g glucose m^-2^). For calculating construction costs, we assumed a growth efficiency of 0.89 (McDermitt and Loomis, 1981). Because measuring combustion heat is laborious and time-consuming, we measured combustion heat for a subset of 1455 samples (33% of the total plant samples). Similarly, the mineral fraction was measured by combustion (4 h at 500°C) for a subset of 96 samples. The subsets of samples were purposely selected to derive average construction costs for each plant organ at different crop stages **(Supplementary Table S1)**. These average values were subsequently used for estimating ADMcc for all plant samples collected across the seven experiments. We did not attempt to distinguish between organic and inorganic N; instead, we assumed that most of the plant tissue N was in organic forms, which is a reasonable assumption considering that other studies have shown that most of the N in ADM exists in organic compounds (Williams *et al*., 1987).

### 2.3 Measurement of absorbed solar radiation during the growing season

The incident, transmitted and reflected photosynthetically active radiation (PAR_I_, PAR_T_, and PAR_R_, respectively) were measured every second and recorded as a 30-minute average in one or two replicates per treatment at each location and year. Sensors were placed in the field when the soybean crop reached the V2 stage (LAI *≈* 0.3). Each line and quantum sensor used for solar radiation assessments was calibrated by the manufacturer and cross-calibration among sensors was performed every year before and after the soybean growing season. Sensors were cleaned from dust and fallen leaves and petioles twice a week during the crop season.

The PAR_I_ was measured above the canopy using a point quantum sensor facing the sky (LI-190SA, LI-COR, Lincoln, NE). A single-line quantum sensor measured the PAR_T_ (LI-191SA, LI-COR, Lincoln, NE) placed at ground level diagonally between two soybean rows. The PAR_R_, from canopy and soil, was measured at 1.5 m above the canopy with a point quantum sensor facing downwards. Sensor height was periodically adjusted to keep the same distance between the sensor and the top of the canopy. To estimate soil reflectance (PAR_S_), we used unpublished data from previous experiments conducted in the same four fields during the 2015 crop season. These previous experiments included an additional single-line quantum sensor facing downwards, placed at 5 cm above the soil and placed diagonally between two soybean rows. Measured PAR_S_ was used to derive the following relationship between PAR_S_ (as a fraction of PAR_R_) and LAI:

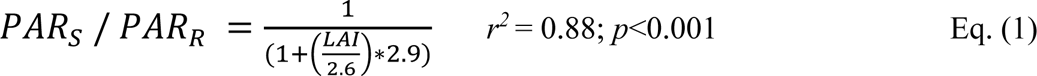

The fitted relationship indicates that the contribution of PAR_S_ to PAR_R_ is largest early in the season, declining afterward as LAI increases. On a seasonal basis, PAR_S_ represents a relatively small fraction of total PAR_I_ (*ca.* 6%). This relationship was subsequently used to estimate daily PAR_S_ for the 2016 and 2017 experiments based on measured LAI and PAR_R_. Finally, the daily APAR was calculated following (Goward and Huemmrich, 1992):

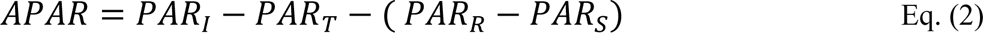

The APAR fraction (*f*APAR) was estimated as the ratio between APAR and PAR_I_. To avoid the inclusion of APAR by non-green LAI during reproductive stages, we corrected APAR using the green-to-total leaf dry matter ratio following (Hanan *et al*., 2002). Likewise, because we could not measure APAR in more than one or two replicates per treatment at each site-year due to logistic and equipment constraints, we estimated daily *f*APAR for the other replicates based on the measured LAI using the relationship reported by Cafaro La Menza *et al*. (2020) derived from the same experimental database (*f*APAR=0.98-exp[-0.54*LAI]; *r^2^* = 0.98; p<0.001). Because LAI was measured weekly, we used a double sigmoidal equation following Fisher *et al*. (2006) to interpolate daily *f*APAR between sampling dates. Finally, daily APAR was estimated by multiplying daily *f*APAR and PAR_I._

### 2.4. Estimation of RUE and data analysis

A logistic model of plant growth from Thornley and France (2007) was fitted to the seasonal trends in cumulative ADM, ADM_cc_, and APAR in each replicate-treatment combination of the seven experiments using GraphPad Prism version 7.0.0 for Windows (GraphPad Software, San Diego, California USA). Analysis of residuals throughout ontogeny was carried out to identify the best-fit model following Hall *et al*. (1995). The RUE (g dry matter MJ^-1^) was estimated as the quotient between cumulative ADM (or ADM_CC_) and APAR between two consecutive ADM samplings, as derived from the fitted logistic models. This approach reduces the experimental error associated with ADM and APAR measurements (Bange *et al*., 1997; Hall *et al*., 1995). Our RUE calculations are based on measured data collected after the V3 stage (LAI*≈* 0.7) to ensure that plants were tall enough so that APAR was not underestimated. Separate estimates of RUE were derived based on accumulated ADM and ADM_CC_; the latter is referred hereafter as RUE_CC_ (g glucose MJ^-1^). Previous studies have shown that RUE may increase up to 8% when root dry matter is included in its calculation (Rochette *et al*., 1995). Because we did not measure root dry matter, we estimated it based on our measurement of ADM and the seasonal pattern in root biomass (as a fraction of the whole plant) reported by Setiyono *et al*. (2010). Subsequently, we re-estimated RUE and RUE_CC_ based on whole-plant dry matter, assuming that the construction cost for belowground dry matter (including roots and nodules) is similar to that of stem dry matter, which is a reasonable assumption based on previously reported data (Amthor *et al*., 1994). Similarly, to account for possible changes in RUE due to seasonal variation in total incoming solar radiation, our whole-plant RUE and RUE_CC_ were adjusted to a total daily incoming solar radiation of 20 MJ m^-2^, based on the relationship reported by van Roekel and Purcell (2014).

Analysis of variance using the PROC GLIMMIX procedure in SAS® version 9.3 (SAS Institute Inc, Cary, NC) was carried out to test the effects of crop ontogeny and N supply on SLN and RUE. A statistical model of repeated measures over time was used with compound symmetry as a covariance structure. The SLN and RUE were tested for normality and homogeneity of variance using PROC UNIVARIATE and no data transformation was needed. The effect of ontogeny was tested by dividing the crop cycle into five phases: V3-R1, R1-R3, R3-R5, R5-R6, and R6-R7. Environment, N treatment (zero and full-N), crop phase, and N by crop phase interaction were considered as fixed effects, whereas replicates were considered random effects. Contrasts were constructed to detect statistically significant differences between N treatments in each crop phase. Relationships between RUE and temperature were assessed using linear regression analysis, separately for each crop phase. Finally, RUE was plotted against SLN and compared against the Sinclair and Horie (1989) theoretical model for understanding how seasonal changes in RUE were associated with variation in SLN as influenced by the N supply. Because the theoretical model was based on RUE estimated from ADM and intercepted total solar radiation, we re-calculated our RUE based on intercepted PAR-based RUE for comparison purposes, assuming that 46% of total incoming solar radiation is PAR_I_ (Sinclair and Muchow, 1999). On average, RUE based on intercepted PAR was 3% lower than that estimated using APAR. Both the theoretical model and our measured RUE were adjusted to a daily incident total solar radiation of 20 MJ m^-2^, which is consistent with the average solar radiation in our experiments. No attempt was made to correct RUE by temperature, diffuse solar radiation, or SLN distribution within the canopy.

## 3. Results

### 3.1. Seasonal patterns in weather, absorbed radiation, biomass, and specific leaf nitrogen

There were no differences in average PAR_I_ and maximum and minimum temperature (Tmax and Tmin, respectively) across environments **(Figure 1)**. In general, solar radiation decreased gradually during the crop season, from *ca.* 12 MJ m^-2^ d^-1^ (V3 stage) to *ca.* 7 MJ m^-2^ d^-1^ (R7 stage). The temperature remained stable (2016) or increased (2017) from V3 until R3, with Tmax and Tmin averaging 29.4 °C and 15.6 °C, respectively. Temperature was lower after R3, averaging 28.2 °C (Tmax) and 15.4 °C (Tmin).

**Figure 1.**
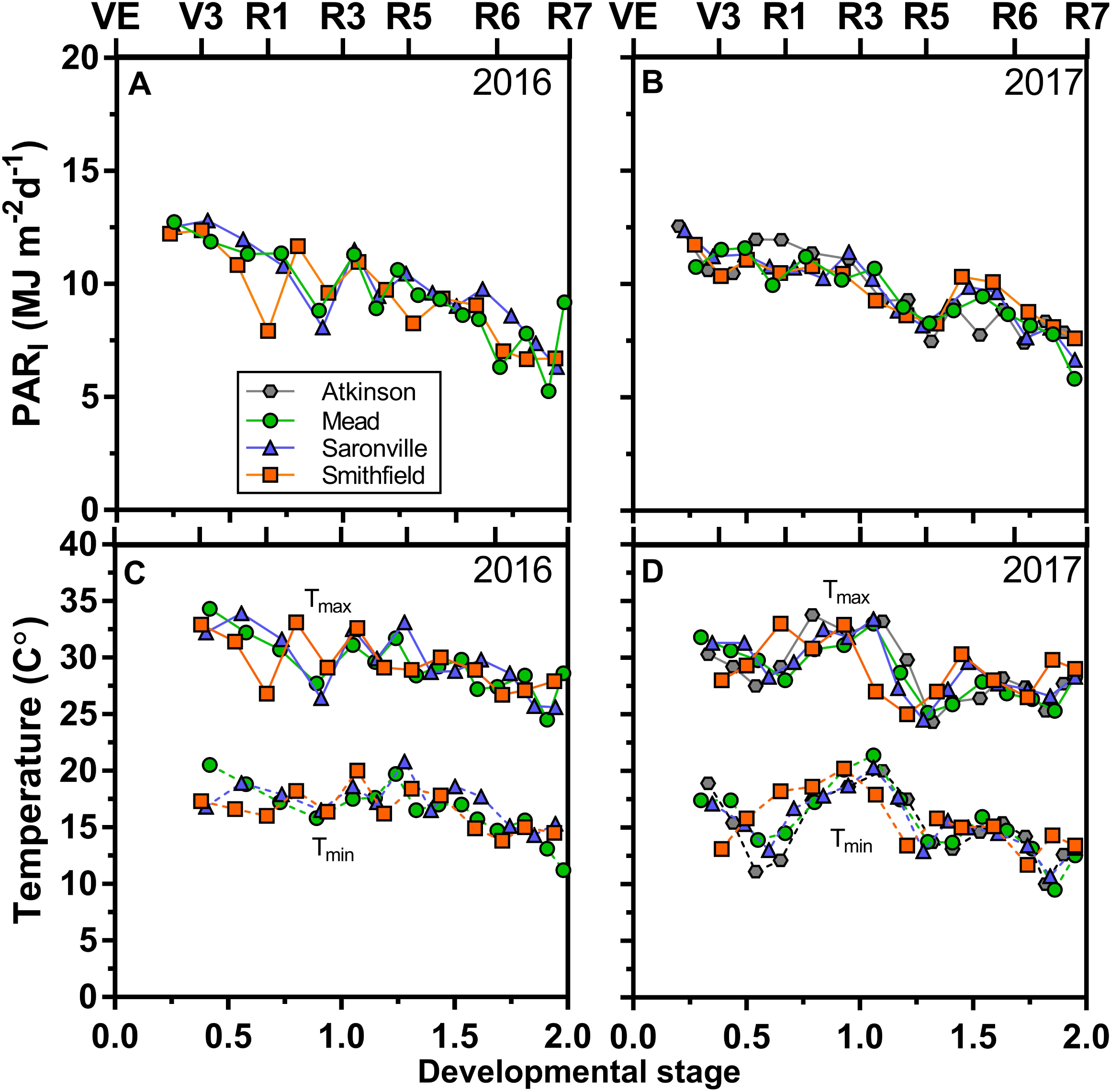
Average (7-d) incident photosynthetically active radiation (PAR_I_)(A and B) and maximum (Tmax) and minimum (Tmin) temperature (C and D) between V3 and R7 stages in seven experiments conducted in Nebraska (USA) at four locations (Atkinson, Mead, Saronville, and Smithfield) during two years: 2016 (A and C) and 2017 (B and D). Phenological time is shown in the bottom x-axis using a dimensionless scale adapted from Lindquist et al. (2005) which allows comparisons to be made among environments. The major development stages defined by the Fehr & Caviness (1977) scale are shown on the upper x-axis; VE: emergence, V3: three fully developed leaves at main stem, R1: beginning of flowering, R3: beginning of pod setting, R5: beginning of seed filling, R6: full seed, and R7: physiological maturity.

The full-N treatment showed slightly but consistently greater *f*APAR between V3 and R5 stages, leading to more ADM accumulation in the full *versus* zero N treatment at the end of the season **(Figure 2)**. There was a significant effect of crop stage, N treatment, and their interaction on SLN **(Table 1)**. Seasonal SLN followed a concave pattern until the middle of the seed-filling phase, followed by an abrupt decline afterward. Early in the season, SLN decreased from 2.5 g N m^-2^ (V3 stage) to 1.5 g N m^-2^ (R1 stage) in both N treatments. After the R1 stage, SLN increased in the full N treatment until reaching a peak around the R6 stage (2.4 g N m^-2^). In contrast, the peak in SLN was less pronounced in the zero N treatment (1.8 g N m^-2^). Shortly after reaching the peak, SLN decreased abruptly in both N treatments, converging into a near-identical SLN at R7 stage (*ca.* 1 g N m^-2^). Seasonal patterns in SLW and leaf N concentration were similar to SLN **(Figure 2)**. However, differences in SLN between N treatments were largely associated with leaf N concentration changes rather than changes in SLW.

**Figure 2.**
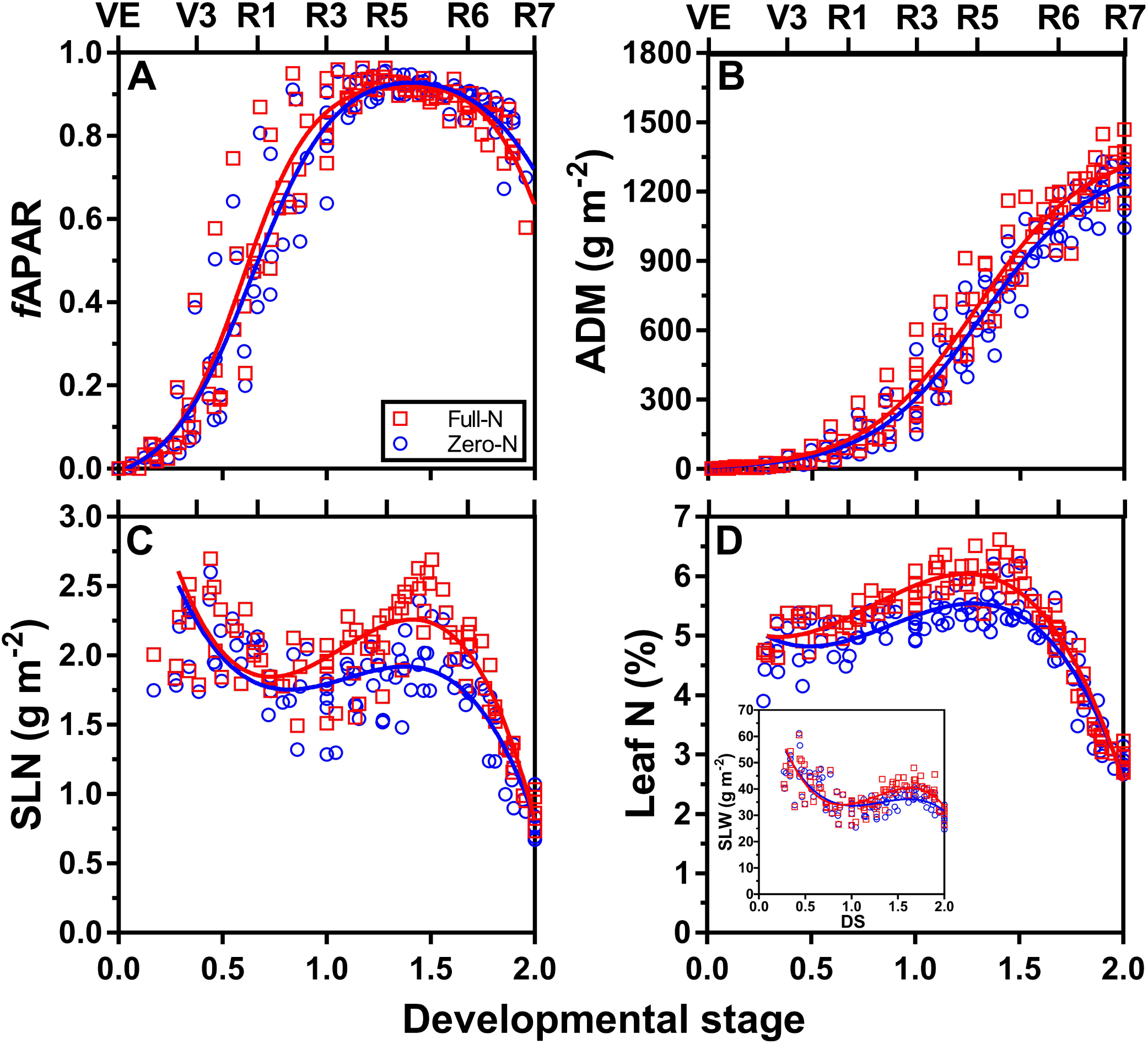
Seasonal trends in (A) fraction of absorbed photosynthetically active radiation (*f*APAR), (B) aboveground dry matter accumulation (ADM), (C) specific leaf nitrogen (SLN), and (D) leaf N concentration in the full-N (red squares) and zero-N (blue circles) treatments as a function of developmental stage which is dimensionless scale adapted from Lindquist et al., (2005) that allows comparisons to be made among environments. The major development stages defined by the Fehr & Caviness (1977) scale are shown on the upper x-axis; VE: emergence, V3: three fully developed leaves at main stem, R1: beginning of flowering, R3: beginning of pod setting, R5: beginning of seed filling, R6: full seed, and R7: physiological maturity. Inset in D shows the seasonal pattern in specific leaf weight (SLW). Solid red and blue lines represent the fitted models for the full-N and zero-N treatment means computed from pooled data across environments.

**Table 1.**
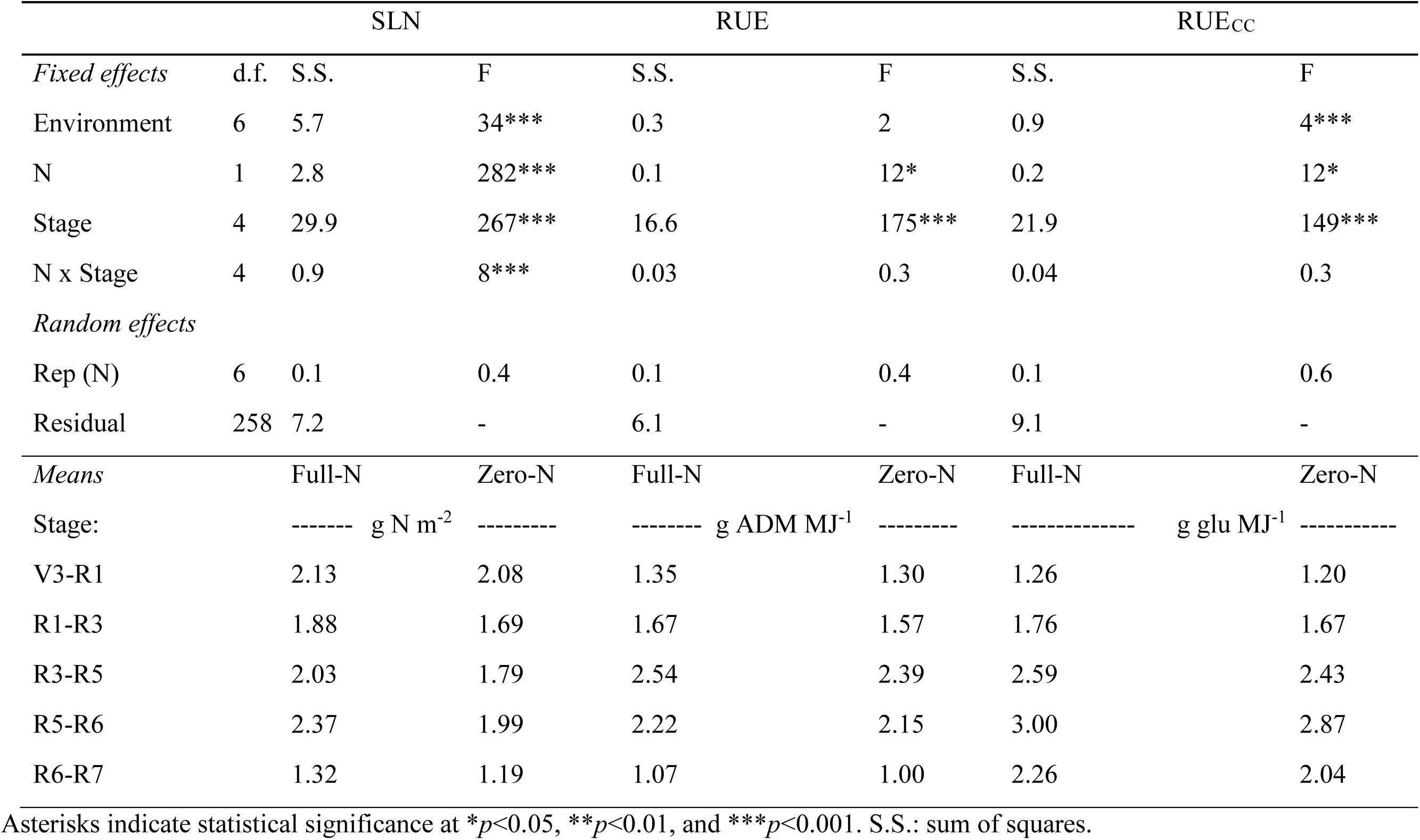
Analysis of variance for effect of nitrogen (N) treatment, crop developmental stage, and their interaction on soybean specific leaf nitrogen (SLN), radiation use efficiency (RUE), and biomass cost-corrected RUE (RUE_CC_). Estimated means for each N treatment at each developmental stage phase are shown.

### 3.2. Variation in RUE as influenced by N supply and temperature

Ontogeny and, to a lesser extent, N supply influenced RUE **(Table 1)**. The seasonal pattern in RUE differed from the pattern in SLN, following a bell-shaped pattern **(Figure 3).** Lowest RUE was observed in early and late crop stages, with a maximum RUE of 2.54 (full N) and 2.39 g ADM MJ^-1^ (zero N) occurring around R5 stage. Higher N supply led to slightly, although consistently, higher RUE in the full *versus* zero N treatment until the early seed-filling phase. Adjustment of RUE by construction costs did not modify the seasonal pattern of RUE, except for (i) a shift in the RUE_CC_ peak to a slightly later R5 stage, and (ii) differences between N treatments that were larger during the late seed-filing phase. Accounting for root biomass and variation in PAR_I_ did not modify the seasonal pattern in RUE **(Figure 3 insets)**.

**Figure 3.**
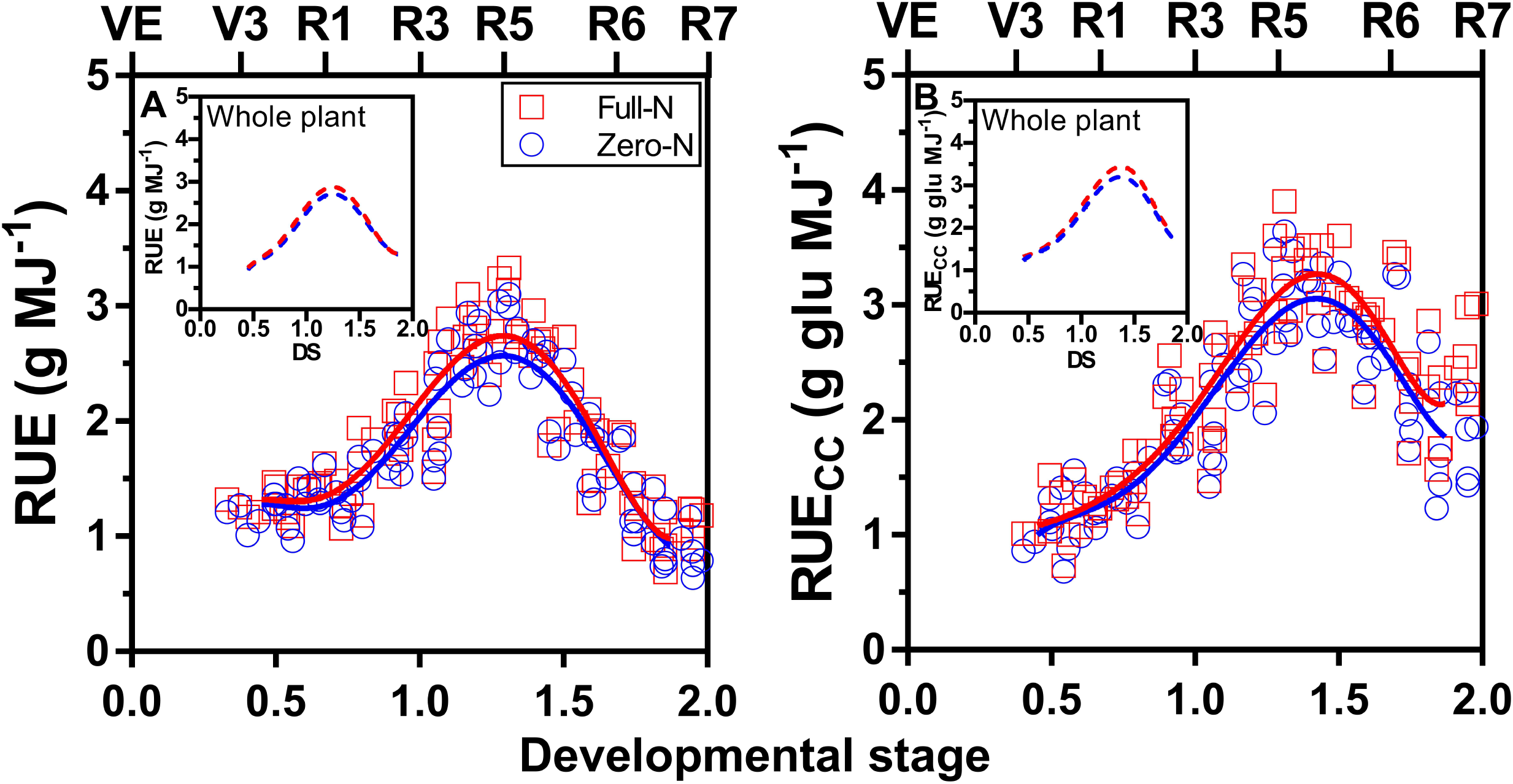
(A) Soybean radiation use efficiency (RUE) in the full-N (red squares) and zero-N (blue circles) treatments as a function of the developmental stage (DS). (B) RUE based on aboveground dry matter adjusted by construction costs (RUE_CC_). The Vn and Rn stages are based on Fehr & Caviness (1977) and shown on the top x-axis. Solid red and blue lines represent the fitted models for the full-N and zero-N treatment means computed from pooled data across environments. Insets show the estimated whole-plant RUE and RUE_CC_, including root dry matter, as a function of the DS, and adjusted to an incident solar radiation of 20 MJ m^-2^ (PAR_I_ = 9.2 MJ m^-2^).

Our analysis also revealed relationships between RUE and Tmax and Tmin; however, these relationships were not consistent among crop stages **(Figure 4)**. For example, there was no relationship between RUE and temperature during the VE-R1, R1-R3, and R5-R6 phases. In contrast, we found a negative relationship between RUE and temperature during the pod setting (R3-R5) and a positive relationship with Tmin during the late seed-filling period (R6-R7).

**Figure 4.**
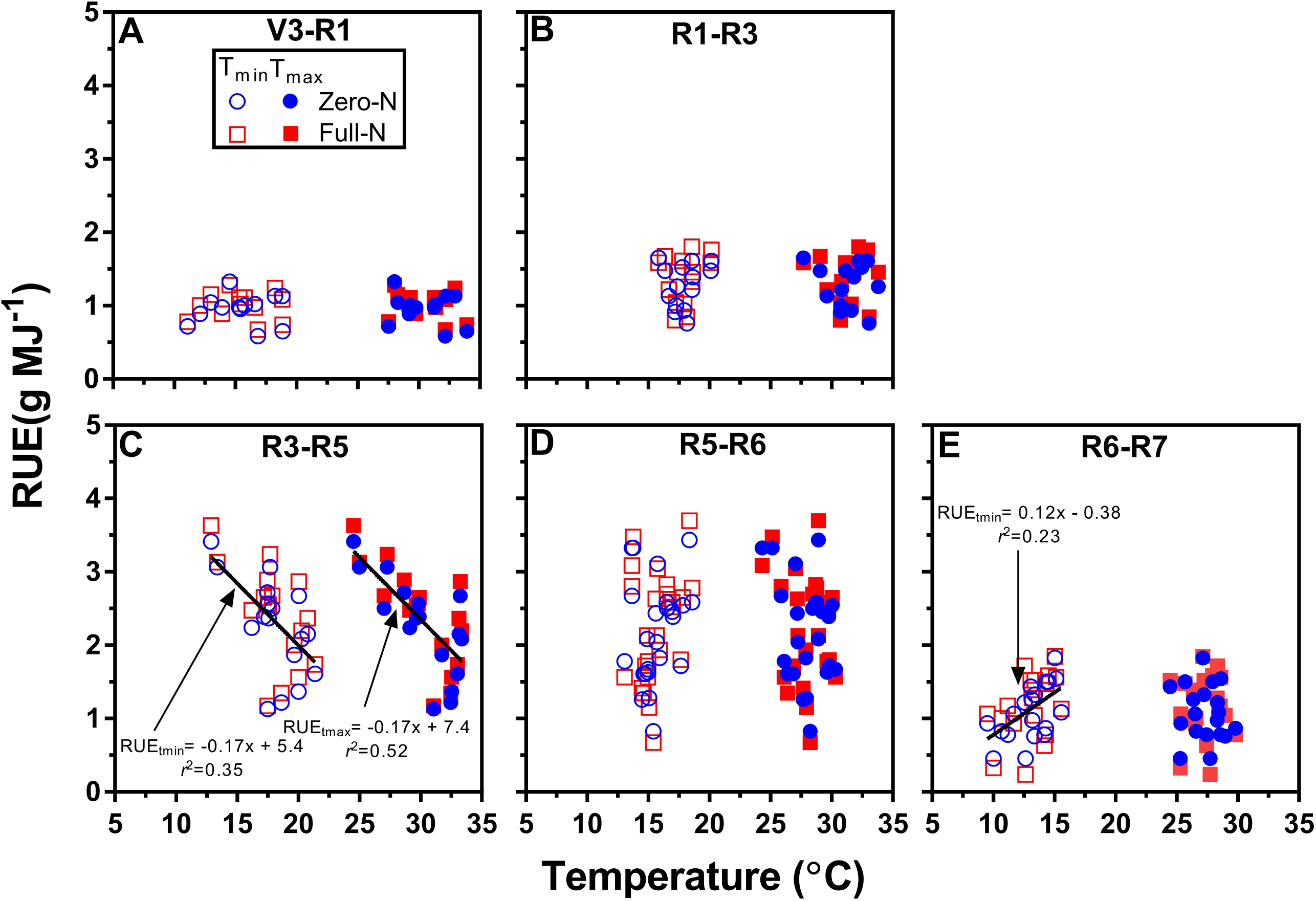
Soybean radiation use efficiency (RUE) as a function of minimum (Tmin) and maximum temperature (Tmax) for different crop phases: VE-R1 (A), R1-R3 (B), R3-R5 (C), R5- R6 (D) R6-R7 (E). RUE was estimated based on aboveground dry matter and absorbed radiation and adjusted to a daily incident solar radiation of 20 MJ m^-2^ (PAR_I_ = 9.2 MJ m^-2^). Fitted linear regression models are shown only when statistically significant (p<0.05); associated parameters and coefficient of determination (r^2^) are shown.

### 3.3. Relationship between RUE and specific leaf nitrogen

Comparing the measured RUE (or RUEcc) and SLN with the theoretical model developed by Sinclair and Horie (1989) indicates that the model reproduces well the sharp decline in RUE driven by SLN during the seed-filling period **(Figure 5a; Supplementary Figure S11)**. Similarly, there was a significant relationship between RUEcc and SLN, but only for growth after R5 stage **(Figure 5b; Supplementary Figure S1)**. The decline in RUE and RUEcc between R5 and R6 (*ca.* -15%) was related to the sharp decrease in SLN (*ca.* -40%). Interestingly, the ample N availability in the full-N treatment did not prevent the decline in SLN and RUE during the seed- filling period.

**Figure 5.**
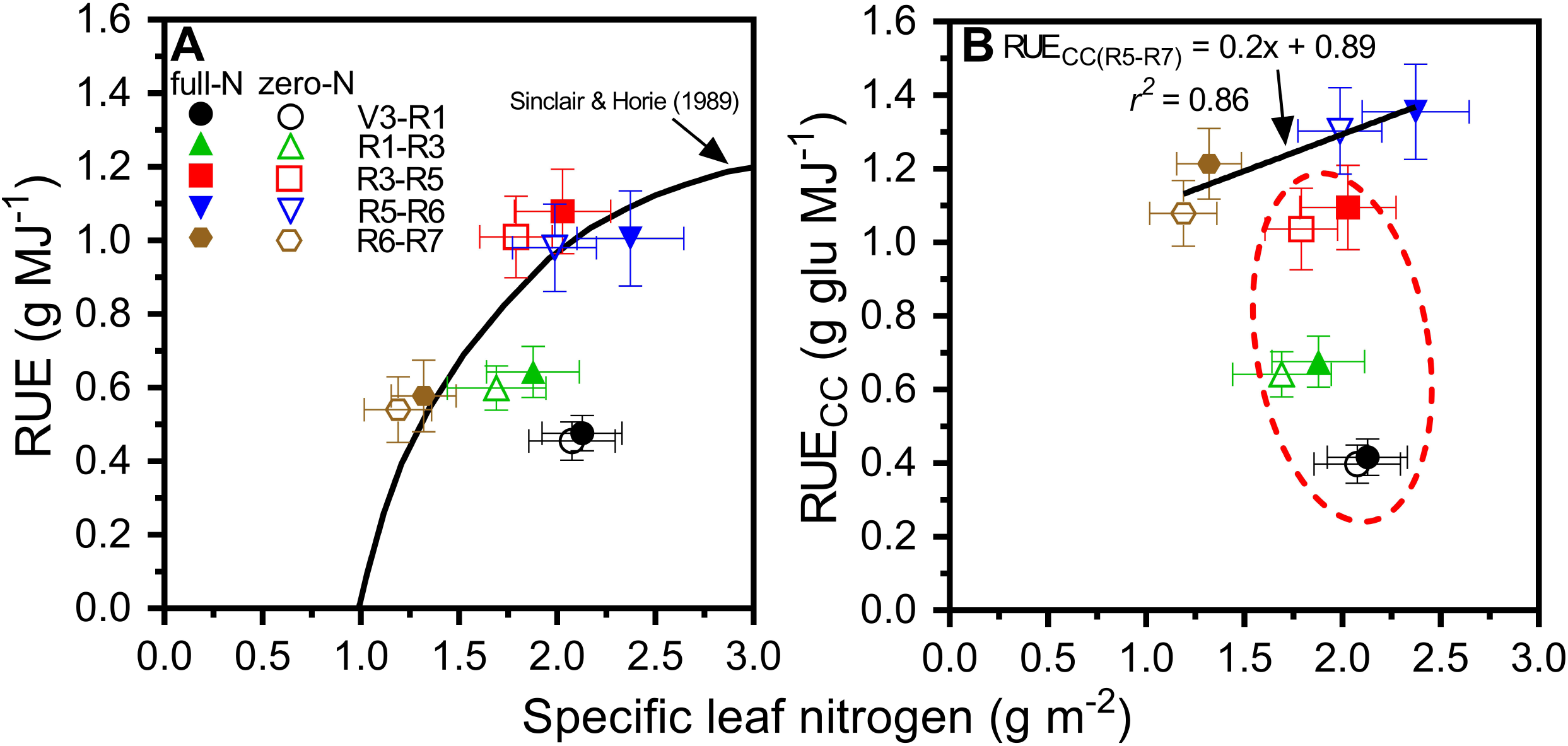
(A)Relationships between radiation use efficiency (RUE) and specific leaf nitrogen (SLN), with RUE estimated based on aboveground dry matter and intercepted solar radiation, and adjusted to a daily incident solar radiation of 20 MJ m^-2^. Also shown is the theoretical model developed by Sinclair and Hoire (1989) relating RUE and SLN (black line). (B) Same as panel (A) but with RUE estimated based on aboveground dry matter after correction by construction costs (RUE_CC_). Also shown in (B) the fitted linear regression between RUE_CC_ and SLN after R5 and associated parameters and coefficient of determination (r^2^). Dashed red line encircle the vegetative and early reproductive stages in which RUE_CC_ significantly chages independently of SLN. In both panels, each data point represents the average RUE (or RUE_CC_) and SLN for the full-N (full symbols) and zero-N (empty symbols) during different crop phases, which are shown with different symbols. Vertical and horizontal lines indicate the standard error of the mean. Relationships based on all data points are shown in Supplementary Figure S1.

The Sinclair and Horie (1989) model did not perform well in predicting RUE patterns during vegetative and early reproductive stages (VE-R3), as there was no relationship between RUEcc and SLN during these crop stages **(Figure 5; Supplementary Figure S1)**. Likewise, the sharp change in RUE observed during the middle of the season (R1-R5) could not be attributed to changes in SLN. For example, average RUE and RUEcc during R3-R5 was 65% greater than that during R1-R3, and this change was accompanied by a modest (+7%) increase in SLN.

The Sinclair and Horie (1989) model did capture reasonably well the increase in RUE due to higher SLN in the full *versus* zero N treatment, although it overestimated the response **(Figure 5)**. According to the model, the difference in SLN between the full and zero N treatments during the R1-R3, R3-R5, and R5-R6 should have led to a concomitant increase in RUE by ca. 14%. In contrast, measured RUE only differed by 8%, 7%, and 3% in the full *versus* zero N treatment during the R1-R3, R3-R5, and R5-R6 stages, respectively.

### 4. Discussion

Seasonal changes in RUE and its underlying drivers were investigated using detailed ADM and APAR data collected during the crop season and considering changes in biomass composition, root dry matter, and incident radiation **(Figure 3)**. The experimental setup helped generate a *wide* range of SLN (0.8-3.0 g m^-2^) to evaluate the importance of SLN in explaining seasonal changes in RUE **(Figure 2)**, and to compare those changes with an existing theoretical model **(Figure 5)**. A sharp decrease in RUE during seed filling was closely related to the decline in SLN, which was consistent with Sinclair and Horie (1989) **(Figure 5)**. However, the theoretical model did not effectively predict the observed RUE changes during the vegetative and early-reproductive stages of growth, even when root biomass and changes in incident radiation were accounted for. Also, the model did not capture the sharp increase in average RUE in the middle of the season (R1-R5). These experimental findings have important implications for modeling crop yield as these phases partly overlap with the critical period for seed number determination in soybean (Monzon *et al*., 2021). Given the uncertainty in predicting RUE based on SLN, our results suggest that early-season RUE can be better estimated based on DS, using a re-scaled version of **Figure 3** (with 0 and 1 corresponding to the lowest and highest RUE, respectively), and using a constant maximum RUE at the R5 stage of 2.5 g MJ^-1^. This simpler approach may provide a more accurate RUE estimation for the vegetative and early-reproductive stages for soybean crops growing in near-optimal conditions. This approach also would overcome the problem associated with assuming a fixed RUE value during the entire crop season for estimating crop growth, yield, and gross primary productivity (GPP) based on remote sensing data and/or crop models.

Our RUE findings are consistent with those reported in previous studies for sunflower and peanut (Bange *et al*., 1997; Hall *et al*., 1995), showing departures between measured RUE and that predicted from the Sinclair and Horie (1989) model. Possible explanations for the low RUE early in the season may include light saturation when leaf area is small (Dold *et al*., 2017; Hall *et al*., 1995; Rochette *et al*., 1995; Trapani *et al*., 1992). Relative to the sharp increase in RUE from R1-R3 to R3-R5 phases **(Figure 5)**, which was not associated with changes in SLN, we speculate that it can be partly explained by increasing demand for assimilates by the growing pods and seeds, which in turn may stimulate photosynthesis (Setter, 1979). Changes in RUE can also be attributed to seasonal changes in the canopy distribution of SLN as reported for other crop species (Grindlay, 1997; Jamieson and Semenov, 2000; Sadras *et al*., 1993). Shiraiwa and Sinclair (1993) attempted to account for the influence of the distribution of SLN in the soybean canopy on RUE, but their measurements were limited to the R2, R5 and R8 stages, and we are not aware of recent reports assessing changes in canopy SLN distribution for modern soybean varieties. We also found that the Sinclair and Horie (1989) model overestimated RUE changes as a result of increased N supply. Likewise, the earlier study by Shiraiwa and Sinclair (1993) did not find any effect of N supply on SLN, while our study showed an increase in SLN due to higher N supply during the R1-R6 phase **(Figure 5)**. A potential explanation is that, under non-limiting N supply, there is an increased synthesis of vegetative storage proteins (VSP) unrelated to the photosynthetic apparatus (Staswick, 1994). Consistent with this hypothesis, changes in SLN were more closely associated with changes in leaf N concentration than SLW **(Figure 2)**. Future work on soybean N dynamics should assess changes in SLN canopy profile and its associated impact on RUE and the role of VSPs in accumulating N that can be mobilized later on to support seed protein accumulation and growth.

Our study also suggests a negative impact of high temperatures during pod setting (R3-R5), and low temperature towards the end of the seed filling (R6-R7) on RUE, which includes part of the critical period (R3-R6) for yield determination **(Figure 4)**. These results are not consistent with a previous study conducted in the U.S. North-central region, showing no relationship between RUE and temperature (Dold *et al*., 2017). We note, however, that this previous assessment was based on monthly average RUE and temperature, and that study covered only a narrow range of temperatures. Previous studies reporting a negative effect of temperature on soybean focused on seed number and size and leaf-level photosynthesis during seed filling (Egli and Wardlaw, 1980; Ergo *et al*., 2018). As far as we are aware, our study is the first to report the negative effects of high and low temperatures on RUE at canopy level. We speculate that this impact may occur *via* reduced photosynthesis and/or sink activity, but further research on this topic is needed since it is relevant for robust simulation of soybean growth and yield in the context of a warming climate.

While we attempted to account for all possible sources of uncertainty regarding the estimation of RUE, our study may be impacted by several potential confounding factors. For example, our estimate of root dry matter is based on a previous model developed by Setiyono *et al*. (2010), which does not account for root:shoot partitioning changes due to contrasting N supply. Biomass partitioning to root may have changed in the full N treatment as a result of ample availability of mineral N in the soil (Cassman *et al*., 1980). However, these changes would have likely led to a very small alteration of RUE, influencing only the RUE estimated for the full-N treatment during early vegetative stages. Likewise, others have documented changes in RUE due to variation in diffuse radiation (Hammer and Wright, 1994; Sinclair *et al*., 1992). For example, lower RUE has been reported for soybean when the ratio between diffuse and total incident solar radiation is < 0.3, with RUE remaining relatively constant above this threshold (Sinclair *et al*., 1992). In the case of our study region in Nebraska, the ratio between diffuse and total incident radiation ranges from 0.6 (May) to 0.4 (September), suggesting that our analysis was not confounded by seasonal changes in diffuse radiation (Kafka and Miller, 2019). Our method for estimating ADM construction costs also relied on a number of assumptions and inherent uncertainty (Griffin, 1994). However, we note that our values are consistent with those reported by previous studies (e.g., Amthor *et al*. (1994); Koester *et al*. (2014)), and also with the 1.3x ratio of seed-to vegetative biomass estimated by Muchow *et al*. (1993a) for soybean. Despite all these sources of uncertainty, our results are robust and the conclusions valid.

## Acknowledgments

We thank the Nebraska Soybean Board for the financial support of this project. We acknowledge Cathleen McFadden, Loren Isom, and Anjeza Erickson at the University of Nebraska-Lincoln (UNL) and Keith Glewen, Todd Whitney, and Amy Timmerman (UNL Extension) for their technical assistance. We also thank Alencar Zanon, Juan Pedro Erasun, Ana Carolina Duarte Rabelo, Mariano Hernandez, and Agustina Diale for their help with field sampling and laboratory measurements. Finally, we are grateful to the four soybean producers in Nebraska who kindly gave us access to their fields to conduct the experiments.

## Abbreviations

ADM: Aboveground dry matter
ADMcc: ADM corrected by construction costs
APAR: Absorbed photosynthetically active radiation
*f*APAR: Fraction of absorbed photosynthetically active radiation
DS: Developmental Stage
N: Nitrogen
PAR_I_: Incident photosynthetically active radiation
PAR_R_: Total (canopy + soil) reflected photosynthetically active radiation
PAR_S_: Soil reflected photosynthetically active radiation
PAR_T_: Transmitted photosynthetically active radiation
RUE: Radiation-use efficiency
RUEcc: Radiation-use efficiency corrected by construction costs
SLN: Specific leaf nitrogen

## Author contributions

NCLM: Conceptualization, Formal Analysis, Investigation, Methodology, Visualization, and Writing – Original Draft Preparation

TJA: Methodology, Resources, Supervision, Writing – Review & Editing

JLL: Methodology, Resources, Supervision, Writing – Review & Editing

JPM: Supervision, Visualization, Writing – Review & Editing

JMHK: Supervision, Resources, Writing – Review & Editing

GG: Supervision, Writing – Review & Editing

DS: Methodology, Resources, Writing – Review & Editing

RH: Formal Analysis, Writing – Review & Editing

JR: Investigation, Writing – Review & Editing

JES: Resources, Supervision, Writing – Review & Editing

PG: Conceptualization, Funding Acquisition, Investigation, Methodology, Project Administration, Resources, Supervision, Visualization, Writing – Review & Editing

## Data availability

Data are available from the authors upon reasonable request.

## Supplementary material for Cafaro La Menza et al.

**Figure S1.**
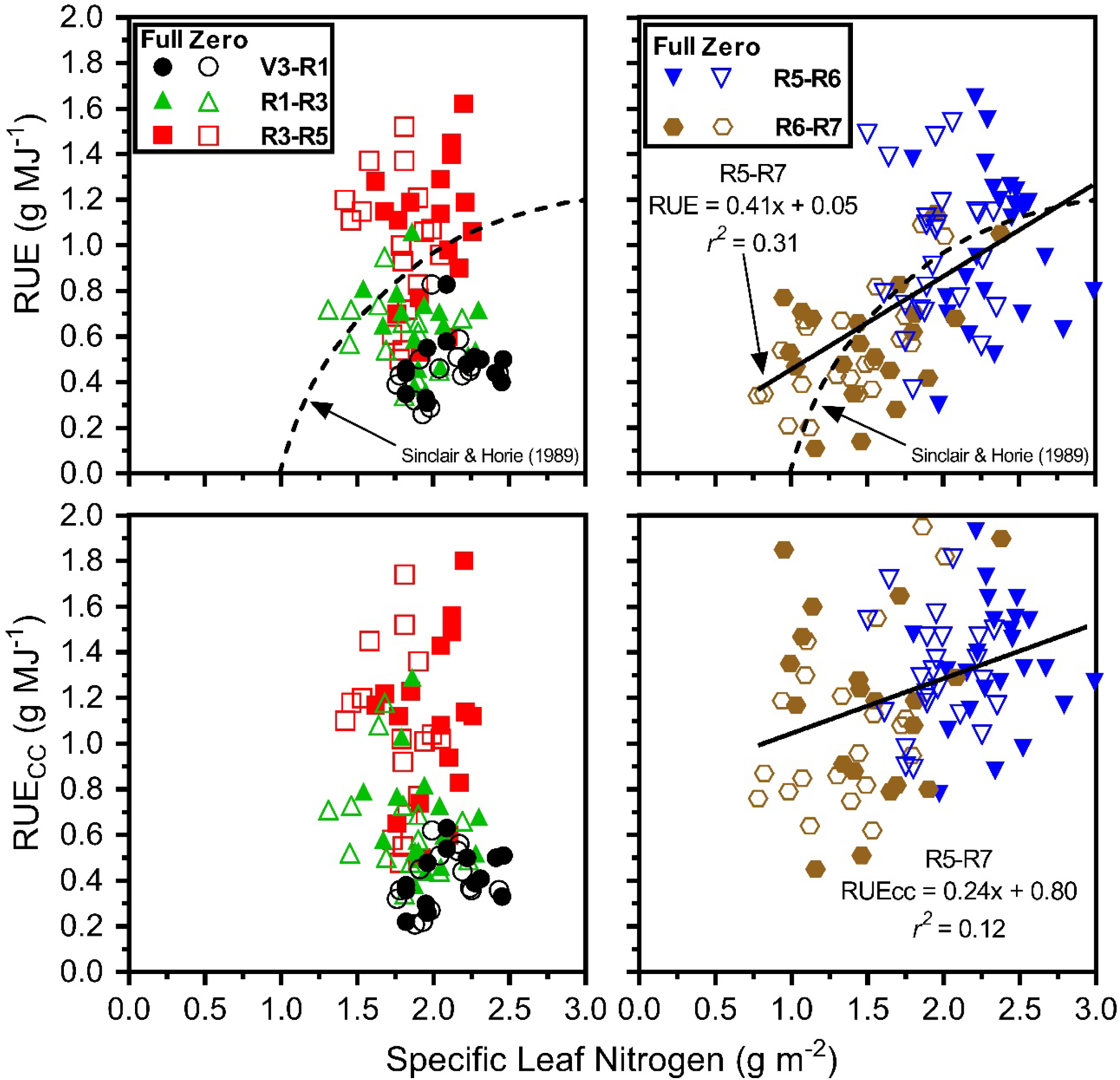
Relationship between radiation-use efficiency (RUE) and RUE estimated based on aboveground dry matter after correction by construction costs (RUE_CC_) as a function of specific leaf nitrogen (SLN) before and after R5 stage (left and right panels, respectively). All RUE values were calculated with intercepted solar radiation and adjusted to an daily incident solar radiation of 20 MJ m^-2^ Also, the fitted linear regression models (p<0.05) and associated parameters and coefficient of determination (r^2^) are shown with a black line, and the theoretical model developed by Sinclair and Hoire (1989) relating RUE and SLN with a dashed line.

**Table S1.**
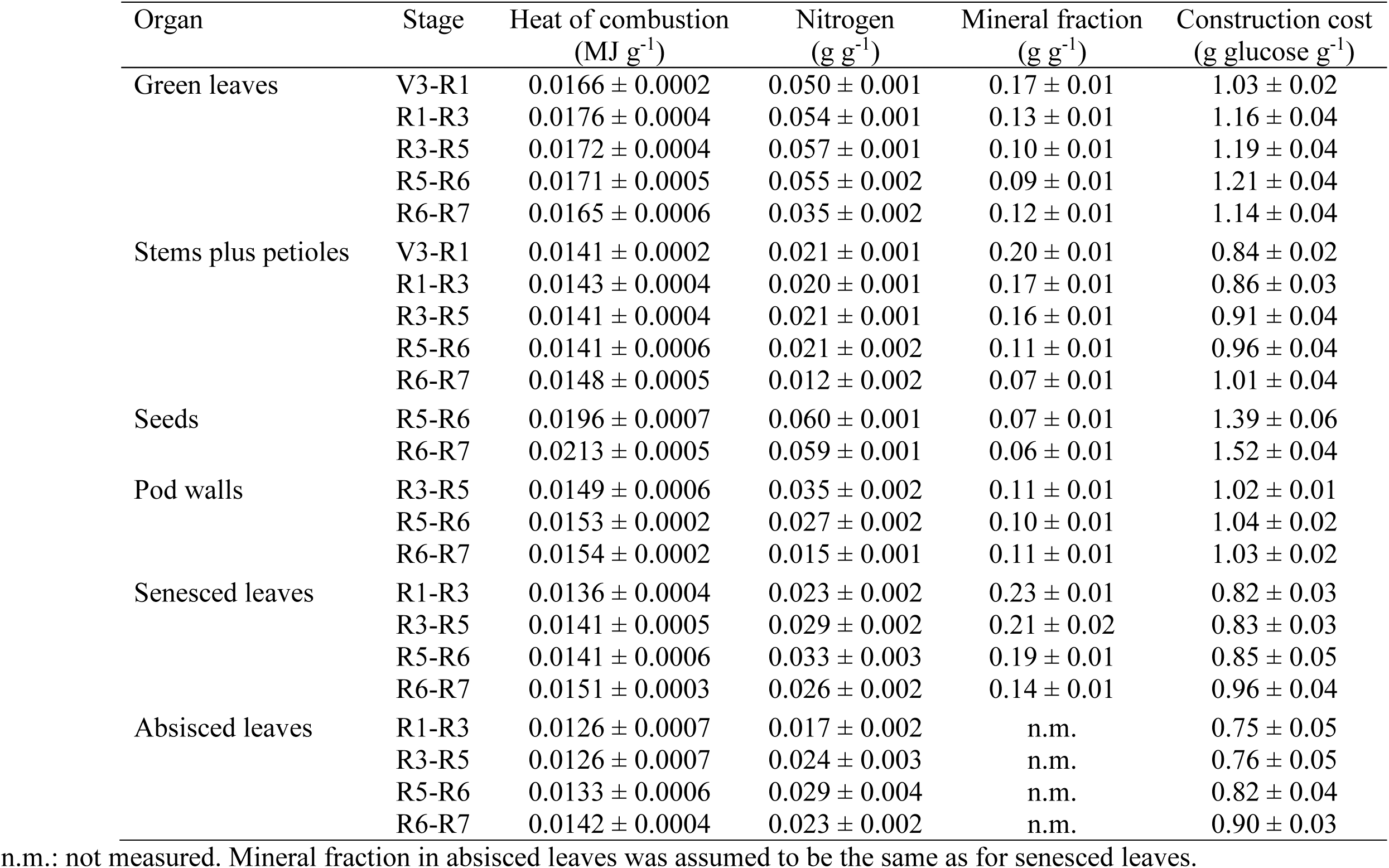
Mean (± standard error) heat of combustion, nitrogen (N) concentration, mineral fraction, and biomass construction cost for different plant organs during different crop stages expressed as the average of full and zero N treatments.

## Footnotes

1 In this manuscript, we used the phenological stages as defined by Fehr and Caviness, (1977) where the Vn indicates vegetative stage with the n indicating the number of fully develop leaves; R1: (beginning bloom) one open flower at any node on the main stem; R3: (beginning of pod) pod 5 mm long at one of the four uppermost nodes on the main stem with a fully developed leaf; R4: (full pod) pod 2 cm long at one of the four uppermost nodes on the main stem with a fully developed leaf; R5: (beginning seed) seed 3 mm long in a pod at one of the four uppermost nodes on the main stem with a fully developed leaf; R6: (full seed) pod containing a green seed that fills the pod cavity at one of the four uppermost nodes on the main stem with a fully developed leaf; R7: (beginning maturity) one normal pod on the main stem that has reached its mature color.

